# Caloric restriction drives age-dependent and integrated remodeling of RNA processing and lipid composition in mouse brain

**DOI:** 10.64898/2026.09.18.752764

**Authors:** Spencer A. Tye, Josef P. Clark, Andrew J. Engeler, Anshit Singh, Rozalyn M. Anderson, Timothy W. Rhoads

## Abstract

Caloric restriction (CR) extends lifespan and delays the onset of age-related diseases. Prior work has established numerous pathways that respond to CR, including growth signaling, metabolism, and inflammatory pathways. However, the molecular mechanisms that connect these pathways to enhanced lifespan and reduced disease risk remain unclear. Here, we demonstrate that RNA processing is altered in brains from mice on CR across adulthood and is linked to changes in lipid composition. Differential gene expression changes at each time point share a high degree of identity and functional overlap, with pathway enrichment including expected changes in metabolic pathways and neurotransmission related pathways. Differential splicing events were observed across adulthood; however, in contrast to the stable transcriptional program, RNA processing changes were dynamic and largely unique to each age group, suggesting regulatory mechanisms that were context specific. CR-induced changes in cortical lipid abundance profiles were highly coordinated across age groups, including increased abundance for lipids with higher degrees of unsaturation. Integrative analysis identified associations between specific lipid classes and transcripts, as well as particular categories of alternative splicing events. These data show that alternative RNA processing is a key mechanism harnessed by CR and linked to lipid homeostasis, providing new insight into metabolic reprogramming in the CR brain.

## Introduction

Aging is a complex process defined by the progressive loss of physiological function and coincident increase in the risk for chronic diseases, including cancer, diabetes, cardiovascular dysfunction, and neurodegeneration (López-Otín et al., 2023). Caloric restriction (CR), without malnutrition, is a dietary intervention that delays aging and the onset of age-related diseases (Longo and Anderson, 2022). CR engages a variety of mechanisms including changes in growth signaling, inflammatory pathways, and metabolic reprogramming, with the molecular details dependent on organism, tissue context, and genetic interaction (Green et al., 2022). However, precise details as to how CR conveys enhanced lifespan still remain unclear. Recent work has uncovered altered transcriptional processing as a prominent feature of CR and possibly a requirement for its full benefits – specific splicing factors are necessary to realize maximal lifespan in long-lived *C. elegans* dietary restriction (DR) models (Heintz et al., 2017; Seo et al., 2016), and changes in transcript exon usage have been identified in multiple tissues from non-human primates on CR (Clark et al., 2025; Rhoads et al., 2018).

RNA processing is the complex process of producing a mature mRNA product, encompassing splicing, capping, polyadenylation, and nuclear export. This involves coordinated handoffs between multiple macromolecular complexes and hundreds of factors all subject to extensive regulation (Carrocci and Neugebauer, 2024). Aging-related alterations to transcriptional processing have been identified in multiple contexts, with dysregulated RNA processing noted as a feature of many age-related diseases (Bhadra et al., 2020). In a relatively small sample group, intron retention events were overrepresented among splicing changes with age in primate cortex (Mazin et al., 2013). Transcriptional elongation rates increase with age in multiple organisms, with interventions that slow RNA polymerase II extending lifespan in fruit flies (Debès et al., 2023). The number of genes with alternative splicing events increases with age in mice undergoing as a result of both normative and premature aging, with the details tissue-and age-specific (Rodríguez et al., 2016). Certain splicing changes are associated with frontotemporal lobar degeneration or Alzheimer’s disease but not found in cognitively normal individuals (Tollervey et al., 2011). Multiple different cancer types are associated with changes in the splicing of cassette exons, with several splicing regulators accounting for a large fraction of changes (Danan-Gotthold et al., 2015). In short, altered RNA processing is substantially associated with aging and the development of age-related diseases but the mechanistic connection with lifespan is as yet poorly defined.

Not surprisingly, nutrient intake and signaling also substantially alter RNA processing. In yeast, resistance to nutrient depletion is dependent on the splicing of specific introns influenced by spliceosome stoichiometry (Parenteau et al., 2025). In *C. elegans*, mTORC1 mediates feeding-regulated splicing of approximately 10% of the genes with multiple isoforms independently of its canonical target S6K (Ogawa et al., 2024). The regulation of glucose-6-phosphate dehydrogenase expression by fatty acids in rat hepatocytes is dependent on a cis-acting exonic element that can silence or enhance exon splicing (Szeszel-Fedorowicz et al., 2006). Alternatively spliced transcripts of fatty acid desaturase enzymes can be regulated by dietary fatty acid levels (Wijendran et al., 2013). Changes to splicing machinery also have downstream consequences for metabolic function, as seen with the knockdown of the splicing factor LUC7L2 that shifts cells towards oxidative phosphorylation (Jourdain et al., 2021). These examples highlight the important role for transcriptional processing in adapting physiology and metabolism.

The degree to which nutrient-driven and age-driven transcriptional processing changes intersect in a model such as CR is still unclear. Here, we examine the effect of CR on transcriptional processing at three different ages in murine brain cortical tissue. We show that despite homeostatic shifts in gene expression machinery, the transcriptional program is largely stable with age. In contrast, CR directed changes in RNA processing appear to be tailored to age, with statistically significant differences in splice events in response to diet in each age group. Next, we used shotgun lipidomics to examine cortical lipid profiles, finding age-related trends in overall unsaturation levels and diet-responsive changes in multiple lipid classes. Finally, we used integrative approaches to identify relationships between gene expression level changes, alternative splicing events, and lipid composition changes, implicating lipid-RNA processing coordination as a regulatory mechanism engaged with CR.

## Results

### CR induces consistent gene expression changes regardless of age

A previously established cross-sectional cohort of B6C3F1/J male mice involved control mice maintained on 95% ad libitum caloric intake and calorically restricted mice maintained on 20% CR (Miller et al., 2017). Animals were placed on the diet at 2 months of age and maintained until sacrifice and tissue harvest at each of three ages: 10 months, 20 months, and 30 months of age. Except where noted, a total of 5 mice per diet per age group were used for all experiments described herein.

We performed RNA sequencing to profile gene expression changes in response to age and diet. CR induced widespread shifts in gene expression in the brains of CR mice compared to control-fed counterparts across all ages examined (**Fig.1A**). These changes were largely consistent regardless of the age of the animals. Principal Component Analysis (PCA) revealed a clear separation by diet along the first principal component as well as a relatively tight clustering of the CR samples compared to the controls (**Fig.1B**). Analysis of differentially expressed genes (DEGs) responsive to CR revealed extensively altered gene expression profiles in response to diet: 2,515, 2,516, and 2,255 DEGs in 10 month-, 20 month-, or 30 month-old animals, respectively (**Fig.1C**) (Table S1). Although the overall pattern is similar regardless of age, the distribution of DEG fold changes in each age group shifted slightly by age. The proportion of upregulated DEGs was highest for 10-month old animals, with 61% of DEGs upregulated. This proportion shifted towards an increasing number of downregulated DEGs for both 20-month (51% upregulated) and 30-month (47% upregulated) animals such that the majority of DEGs are downregulated in 30-month old animals (**Fig.1C**). Despite this trend, the identity of diet-responsive DEGs across all age groups showed a high degree of overlap, with 20% of statistically significant transcripts the same at all ages and approximately half of all DEGs shared between at least two age groups (**Fig.1D**). Functional overlap was also high, with 40% of enriched pathways in common across age groups (**Fig.1D**). In general, direction of effect of constituent genes was also consistent. For example, 2,413 of the 2,515 (96%) statistically significant genes in response to CR have the same direction of effect in 30-month animals, with 1,061 (42%) of those statistically significant with the same direction of effect in both age groups. Thus, the transcriptional program in response to CR is remarkably constant with age.

**Fig. 1.**
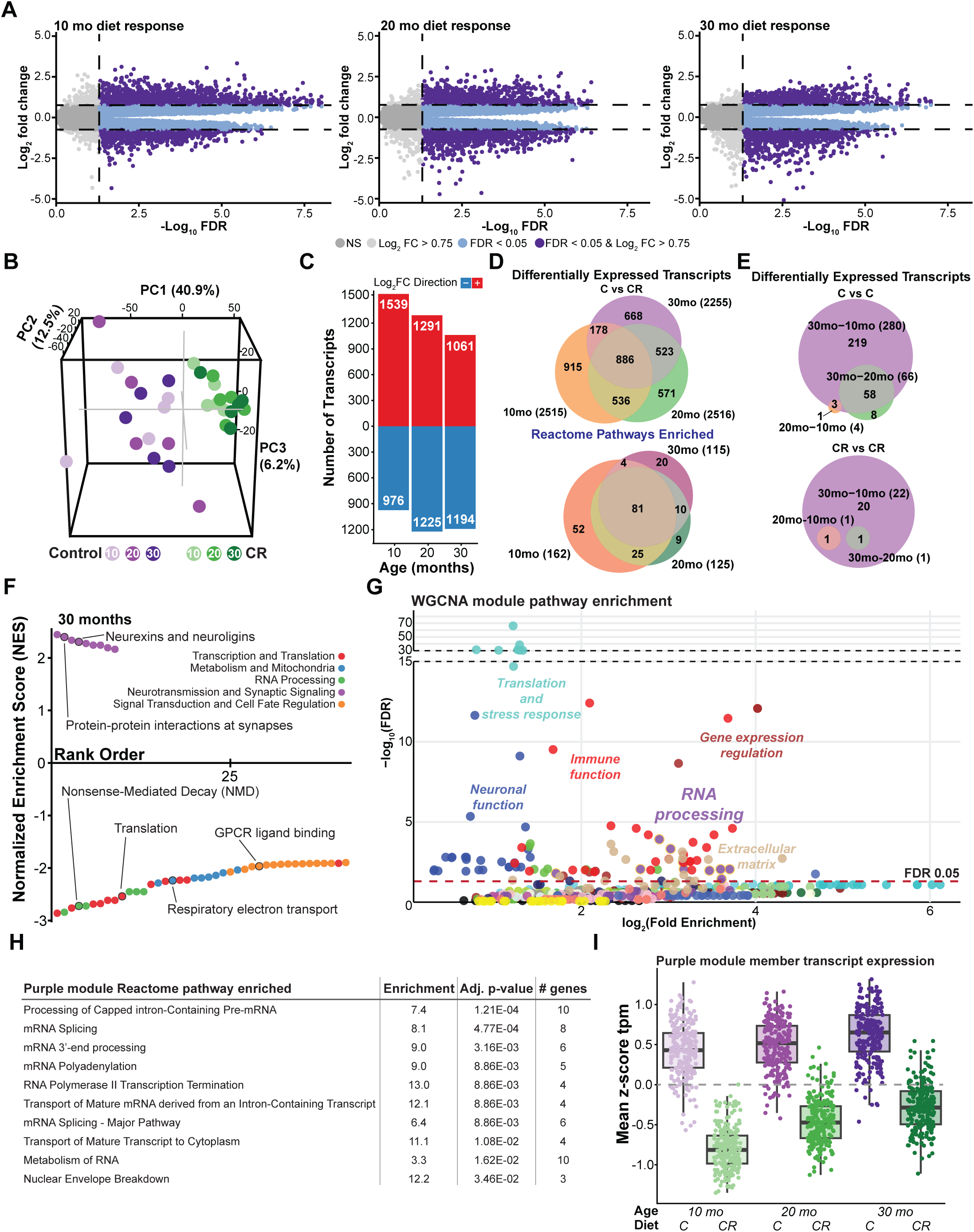
Transcriptomic analysis reveals a consistent response to CR. (**A**) Volcano plots depicting differentially expressed genes (DEGs) in response to diet. DEGs were defined as FDR < 0.05 and |log_2_ fold change| > 0.75. Sample size: Control: N = 5; CR: N = 5 for each age. (**B**) Three-dimensional principal component analysis of the transcriptome (PC1, 40.9%; PC2, 12.5%; PC3, 6.2%), with samples colored by diet and age (control and CR at 10, 20, and 30 months). (**C**) Number of DEGs per age, split by direction of change (upregulated, red; downregulated, blue). (**D**) Euler diagrams of the overlap across ages of diet-responsive DEGs (top) and of significantly enriched Reactome pathways (bottom; FDR < 0.05). (**E**) Euler diagrams of the age-comparison DEGs within each diet - control-versus-control (top) and restricted-versus-restricted (bottom) - across the three ages. (**F**) GSEA using Reactome pathways of the 30-month diet response: normalized enrichment score versus pathway rank, with pathways colored by functional category (transcription and translation; metabolism and mitochondria; RNA processing; neurotransmission and synaptic signaling; signal transduction and cell-fate regulation). (**G**) Gene co-expression WGCNA with per-module Reactome over-representation analysis: each point is an enriched pathway, plotted as log_2_ fold enrichment versus -log_10_ FDR and grouped by the predominant biological function identified within a given module (neuronal function, gene-expression regulation, RNA processing, immune function, extracellular matrix, translation and stress response); the dashed line marks FDR = 0.05. (**H**) Top Reactome pathway enrichments of the purple (RNA-processing) module. (**I**) Mean z-scored expression (TPM) of the purple-module transcripts by age and diet (control, restricted); box = median and interquartile range, points = individual transcripts.

Examining the gene expression changes in response to age alone, we found very few statistically significant transcripts regardless of the ages being compared, with the largest number identified when comparing the 30-month control-fed samples to the 10-month control-fed samples (**Fig. 1E**). The modest effect of age potentially reflects population heterogeneity increasing with age, the effect size in response to age being smaller than in response to diet, and other contributing factors. Of note, only 22 transcripts showed significant expression changes when comparing different age groups of CR mice (**Fig. 1E**), providing further evidence of the consistency of the transcriptional response to CR we observed across ages.

We performed Gene Set Enrichment Analysis (GSEA) to identify the biological functions represented by the diet-responsive gene expression changes. In brain we found the bulk of enriched pathways to represent downregulated transcripts, and pathways typically thought of as activated by CR displayed the opposite behavior. Downregulated pathways include those in categories such as RNA processing, metabolism and energy generation, and transcription and translation, while up-regulated pathways were entirely linked to neurotransmission and synaptic function (**Fig.1F; Fig.S1; Table S1**). The impact of CR opposes the response due to age that has been previously documented for these pathways and the constituent transcripts (Hahn et al., 2023; Souder et al., 2025), suggesting that what we observe in response to CR may reflect the maintenance of a more youthful tissue environment (Zhang et al., 2025). However, other reports indicate increases in some of these pathways and transcripts in response to CR (Tyshkovskiy et al., 2023), highlighting the complexity of these responses and the need for further study. Overall, pathway enrichment profiles were highly similar between age groups moreso than with individual gene expression changes. Approximately 40% of identified enriched pathways were shared among all 3 age groups, with 60% of pathways shared among at least two age groups, suggesting that the transcriptional response to CR at the biological pathway level is highly similar regardless of age or time on diet (**Fig.1D**).

To elucidate more granular detail on the functions associated with the observed gene expression changes, we performed Weighted Gene Co-expression Network Analysis (WGCNA) and used pathway Over Representation Analysis (ORA) to identify biological functions associated with individual modules. Of the 23 modules identified by WGCNA, nine had substantial pathway enrichments that passed statistical significance and suggested coherent function, including modules enriched for neuronal function, immune function, gene expression regulation, translation, and extracellular matrix associated pathways (**Fig.1G**; **Table S2**). The use of WGCNA allowed for a robust RNA processing related signal to be more apparent, with the purple module consisting almost exclusively of RNA-processing related pathways (**Fig.1H**). The expression of transcripts within this module was strongly downregulated in response to CR at all ages, with a slight narrowing of the average effect size in the oldest animals (**Fig.1I**). These findings highlight remodeling of factors associated with RNA processing, consistent with previous studies, and indicate tissue-specificity in the response to CR with the direction of effect opposite (i.e. repressed rather than induced) to that observed in other tissues (Rhoads et al., 2018; Swindell, 2009).

### CR induces a homeostatic shift in gene expression machinery

Given the substantial downregulation observed across factors involved in many aspects of RNA processing, we next asked how the gene expression machinery responded to diet and age. Using a consensus approach, we constructed lists of core and regulatory splicing factors based on the literature (Cvitkovic and Jurica, 2013; Papasaikas et al., 2015). For simplicity, we divided the splicing factors into “core,” or indispensable, and “regulatory,” or facultative, splicing participant categories. Of these 510 factors identified in our data set, approximately 300 were responsive to diet by an FDR less than 0.05 for each age comparison (**Fig.2A**). When considering all factors together there is a tendency towards downregulation in response to CR; however, among the “core” factors specifically, the bias was considerable and the vast majority of the core splicing factors were downregulated (**Fig.2B**). Other parts of the gene expression machinery also displayed a trend towards downregulation, including general transcription factors (**Fig.2C**), mRNA export-associated transcripts, and ribosomal mRNA transcripts (**Fig.2D**) (**Table S3**). This homeostatic downregulation across all ages would seem to suggest an overall reduction of gene expression in response to CR at all ages. However, this does not align with the transcriptional response we observe, where the number of factors downregulated is only the majority at 30 months of age, and the average fold change across statistically significant transcripts is negative also only at 30 months of age (10 mo: 0.27; 20 mo: 0, 30 mo: -0.17). Gene expression regulation by CR is thus clearly dynamic and subject to input from multiple regulatory layers such that gene expression readouts do not tell the whole story.

**Fig. 2.**
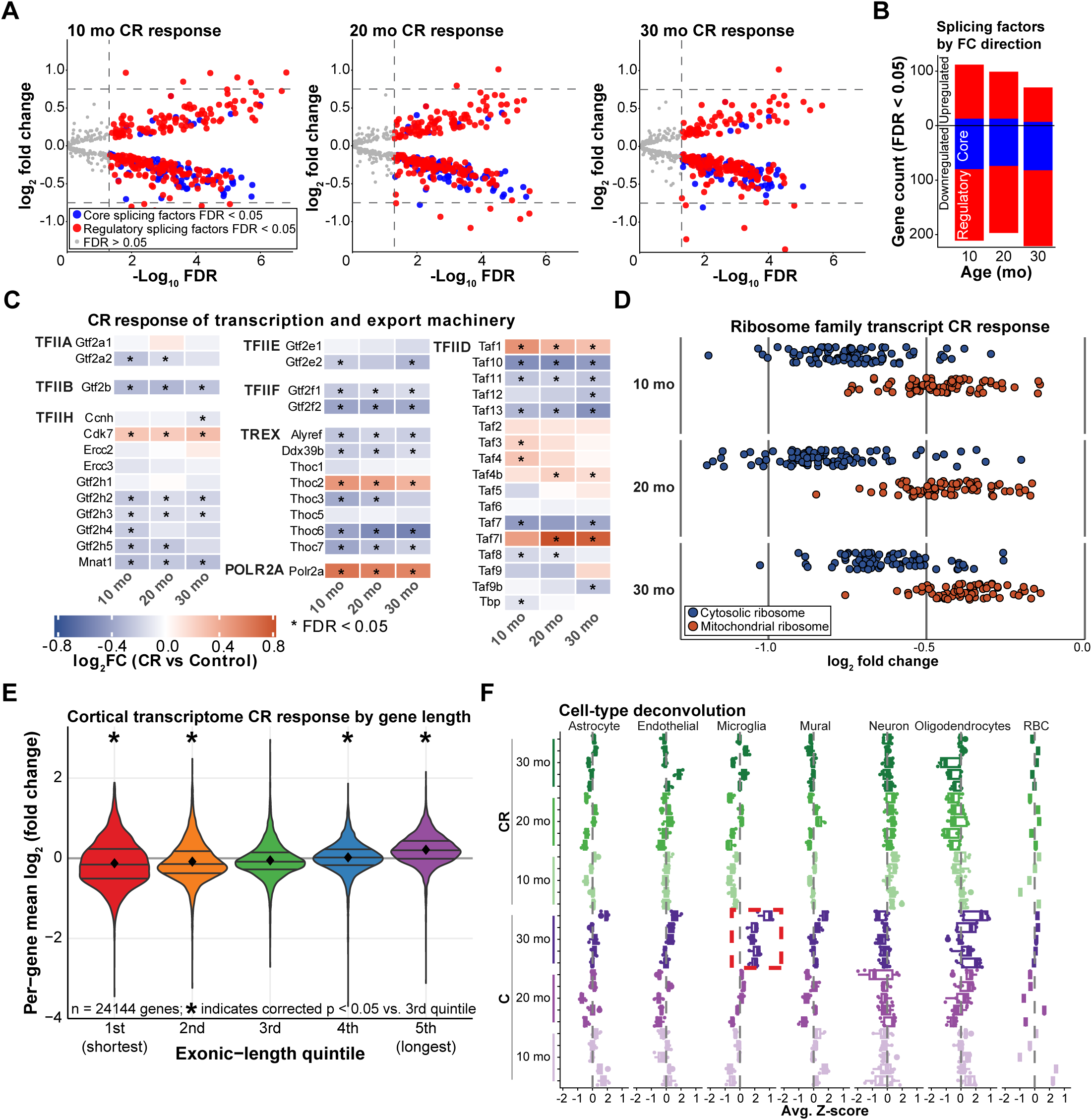
Caloric restriction remodels gene expression machinery. (**A**) Volcano plot depicting the diet response of factors associated with the splicing machinery at 10, 20, and 30 months. Points are colored by class - core splicing factors (blue) and regulatory splicing factors (red) at FDR < 0.05, with non-significant genes (FDR > 0.05) in grey; dashed lines mark FDR = 0.05 and the log_2_ fold-change cutoff (|log_2_ fold change| > 0.75). (**B**) Number of significant (FDR < 0.05) splicing factors by age, split by class (core, regulatory) and by direction (up versus downregulated). (**C**) Diet response of the transcription-initiation and mRNA-export machinery, grouped by complex (TFIIA, TFIIB, TFIID, TFIIE, TFIIF, TFIIH, POLR2A, TREX); columns are age (10, 20, and 30 months) and asterisks mark FDR < 0.05. (**D**) Diet response of ribosomal-protein encoding transcripts by age, colored by cytosolic versus mitochondrial ribosome. (**E**) Per-gene mean log_2_ fold change binned by exonic-length quintile (1st, shortest, to 5th, longest), across n = 24,144 genes; violins show the distribution and the diamond marks the mean. (**F**) Cell-type deconvolution: estimated cell-type signature scores (average z-score) for each brain cell type (astrocyte, endothelial, microglia, mural, neuron, oligodendrocyte, RBC) by diet (restricted, control) and age (10, 20, and 30 months).

We wanted to also determine if there were other aspects of the transcriptome that could influence the changes we observed. We first examined whether there was a length bias observable across the expression changes in response to CR. Increased age leads to a decline in the expression of longer transcripts and thus a shorter transcriptome (Stoeger et al., 2022). We divided the transcriptome into full exonic-length quintiles, finding longer transcripts are upregulated compared to shorter ones (**Fig.2E**), consistent with CR counteracting an age-associated decline in expression for very long transcripts. We also examined the potential contribution from changes in cell type composition by performing RNAseq deconvolution via the *BrainInABlender* R tool (Hagenauer et al., 2018). We observed a mostly consistent cell-type population across age and diet, suggesting that bulk gene expression changes observed are unlikely to be due to shifting cell-type distributions (**Fig.2F; Table S4**); however, expression for microglial markers (red box) in the oldest control animals was increased, a phenomenon that was one of the stand-out observations from large-scale single-cell brain aging data (Hahn et al., 2023). The increase was not observed among the CR animals, suggesting that CR maintains a more youthful cellular environment. Overall, these data highlight age-associated transcriptome changes that are not present in animals undergoing CR, while indicating additional layers of regulation contribute to gene expression changes in response to CR.

### CR harnesses RNA processing in an age-specific manner

To uncover more detail on the changes to RNA processing influenced by CR, we next asked whether individual alternative splicing events (ASEs) were altered by diet. Using the tool SpliceWiz (Wong et al., 2024), we identified numerous statistically significant ASEs, detecting 86, 55, and 89 events in the 10-, 20-, and 30-month old CR mice, respectively, when compared to age-matched controls (**Fig. 3A; Table S5**). These events included alternative 3’ splice sites (A3SS), alternative 5’ splice sites (A5SS), alternative first exons (AFE), alternative last exons (ALE), intron retention (IR), and skipped exons (SE). A3SS was the most common category of significant event type in response to diet across age groups, accounting for 57% of all statistically significant alternative splicing events in response to diet regardless of age. A3SS events were also the substantial majority of the significant events within each of the 10- and 20-month age groups (**Fig.3B**). However, the distribution of significant event types in response to diet in the 30-month old animals varied considerably. For example, AFE and IR events were substantially more numerous, constituting 28% and 24% of the significant diet-responsive events in the 30-month group. This suggests that unlike the overall profile of gene expression and corresponding biological pathway responses to CR, which were generally similar regardless of animal age, changes to alternative splicing were more specific and related to the cellular context at each age. Corroborating this, the overlap of the identity of the individual events between the age groups was generally poor (**Fig.3C**), with only 16% of events being shared between at least two age groups and only four events in common to all three. The overlap is marginally improved when considering the genes in which the significant alternative splicing events are occurring (**Fig.3D**). Because of the expression downregulation observed for ribosomal mRNAs, we examined reads for the individual IR ASE in *Rpl37a* via a sashimi plot (**Fig.3E**), which shows a reduced relative usage of the exon-exon junction and an increase in coverage of the intervening intron in CR samples. The IR event in question is predicted to be a nonsense-mediated decay candidate, a potential mechanistic explanation for expression downregulation.

**Fig. 3.**
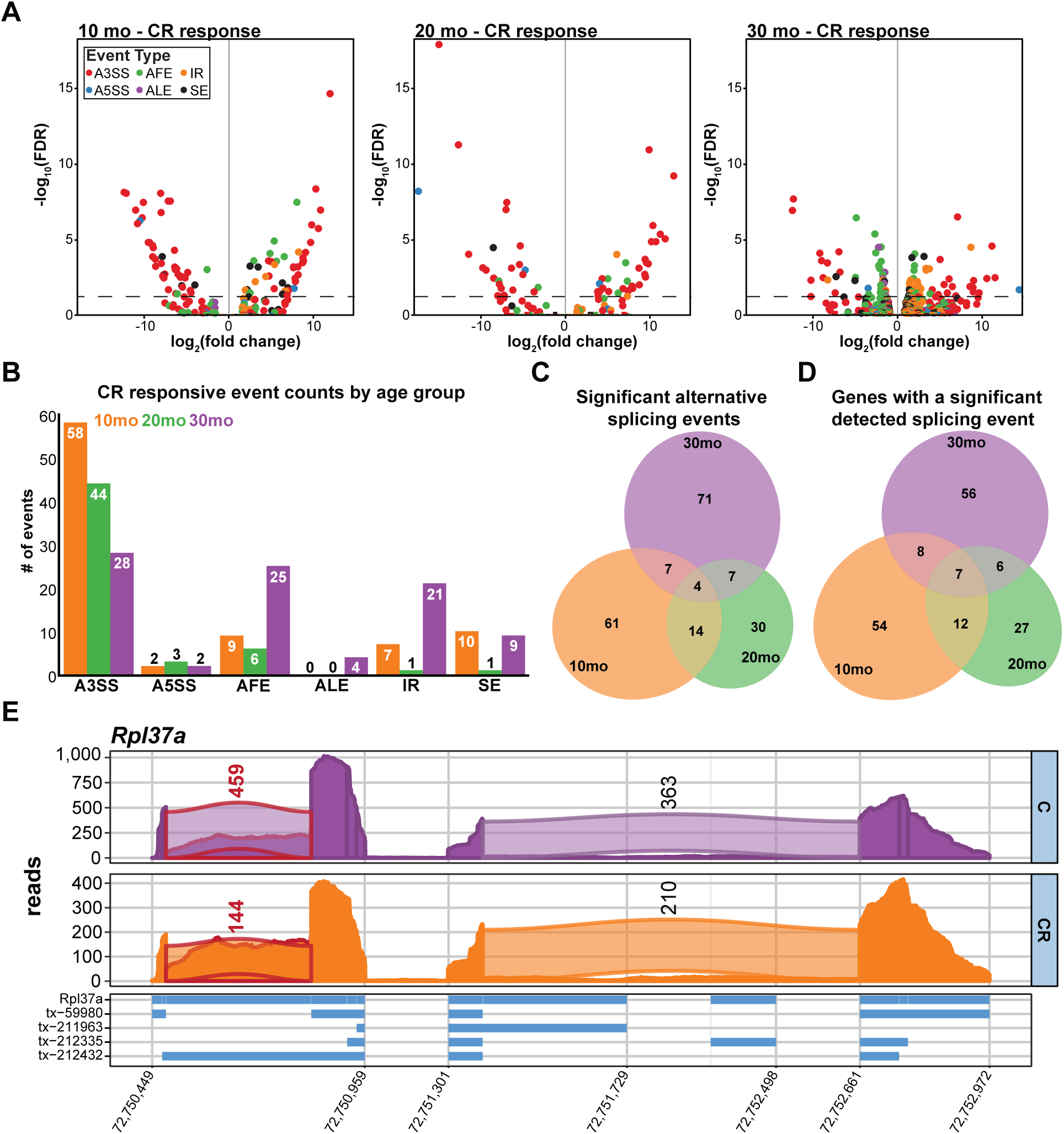
Alternative splicing in response to CR is age-specific. (**A**) Volcano plots depicting differential alternative-splicing events in response to diet at 10, 20, and 30 months; the dashed line marks FDR = 0.05, and points are colored by event type (A3SS, A5SS, AFE, ALE, IR, SE). (**B**) Number of significant (FDR < 0.05) diet-responsive events per event type, grouped by age. (**C**) Euler diagram of the significant (FDR < 0.05) events shared among the three ages. (**D**) Euler diagram of the genes carrying at least one significant (FDR < 0.05) event, shared among the three ages. (**E**) Sashimi plot of Rpl37a, displaying read coverage across the locus in control (C) and restricted (CR) diets, with splice-junction read counts and transcript-isoform tracks below. Reads that contribute to the differential event identification are highlighted in red.

We next asked whether coordinated patterns of splicing variation associated with diet and/or age by performing WGCNA using splicing event quantification (PSI) rather than gene expression. A PSI matrix containing a total of 30,000 splicing events that passed filtering criteria and were quantified in all 3 diet-contrasts was used as input, resulting in 45 modules; for simplicity, we focused on the top 20 modules as ranked by maximum module correlation value with one of the traits (**Fig.S2**). This process identified two modules as significantly associated with diet and the diet x age intersection: the red module, which was negatively correlated, and the blue module, which was positively correlated (**Fig.4A**). The event type composition of the input 30,000 events was: 60.7% IR, 14.1% SE, 10.6% AFE, 6.5% A5SS, 6.3% A3SS, 1.3% ALE, and 0.4% MXE (**Fig.4B**, *top*). Comparing the event-type composition of the diet- and age-associated modules with the full input set, we identified an enrichment for IR events within the blue module (Fisher’s exact test FDR 3.72×10^-81^) and a depletion of AFE, A5SS, A3SS, and ALE events (FDRs 9.67×10^-32^, 9.18×10^-19^, 6.35×10^-25^, and 1.21×10^-4^, respectively) (**Fig.4B**, *bottom left*). The red module was significantly enriched for AFE, SE, ALE event types and depleted of IR events compared to input (Fisher’s exact test FDRs 8.86×10^-15^, 2.68×10^-5^, 1.6×10^-5^, and 2.21×10^-13^) (**Fig.4B**, *bottom right*) (**Table S6**). Within the blue module, we identified a substantial enrichment for ribosomal mRNA transcripts among the population of IR events. The splicing WGCNA input of 30,000 events occurred in 8,686 genes (36% of the total number of genes represented in the transcriptome). There are 162 protein-coding ribosomal genes in mice (KEGG), with 82 present in the WGCNA input and containing an IR event; 51 of those were within the blue module. The majority of these events were concentrated among the cytosolic large ribosome subunit and all but one indicated increased intron retention in response to CR (**Fig.4C**). Increased intron retention has been associated with aging in *Drosophila* and in mouse brain (Adusumalli et al., 2019), although no one clear pathway stands out as preferentially affected. The increased intron retention in response to CR identified here was most evident in ribosomal genes and may represent an adjustment of ribosomal stoichiometry, since IR has been previously implicated in the mechanisms of adaptive ribosomal protein abundance regulation in yeast (Petibon et al., 2016).

**Fig. 4.**
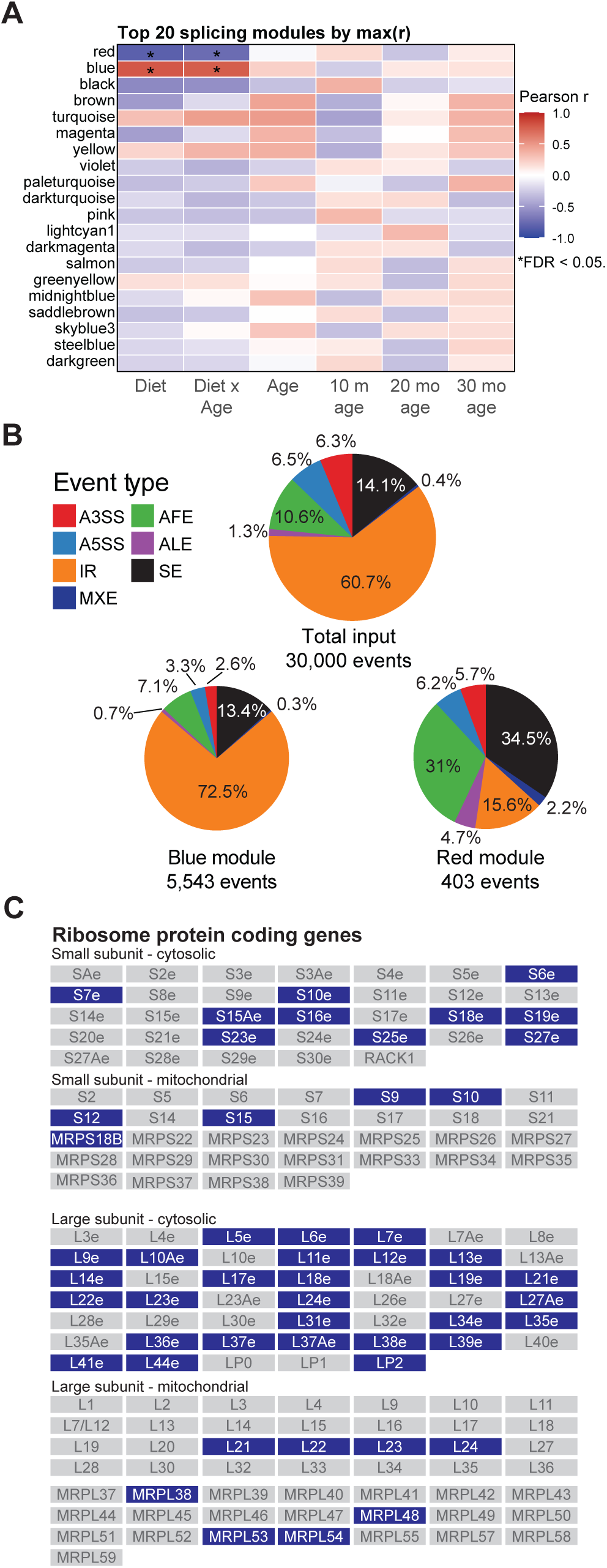
WGCNA analysis of splicing events reveals association of intron retention with the diet response. (**A**) Module-trait correlation heatmap for the top 20 splicing modules (ranked by maximum |r|). Rows are modules (labeled by color); columns are traits: diet, the diet x age interaction, continuous age, and indicators for each age group (10, 20, and 30 months). Cell color is the Pearson correlation between the module eigengene and the trait; asterisks mark correlations significant at FDR < 0.05. (**B**) SpliceWiz event-type composition (A3SS, A5SS, AFE, ALE, IR, MXE, SE) of the full WGCNA input (30,000 events) and of the two diet-associated modules, blue (5,543 events) and red (403 events). (**C**) Atlas of ribosomal-protein genes - cytosolic and mitochondrial small and large subunits - with genes carrying an intron-retention event included in the blue module highlighted.

Collectively, these findings indicate CR remodels RNA processing in the brain in an age-dependent manner, in contrast to changes at the gene expression level that are more strongly diet-driven and age-agnostic. This suggests that transcriptional processing is an important and orthogonal layer of the CR response, and perhaps a mechanism for fine-tuning cellular adaptation to decreased nutrient intake.

### CR drives dynamic reprogramming of brain lipid composition

In identifying the requirement for intact splicing machinery in the lifespan benefits of dietary restriction (DR), Heintz *et al*., identified a lipid catabolism signature among genes showing increased IR with age. DR prevented this increase in all but the oldest animals (Heintz et al., 2017). Lipids are critical to cellular and organismal function, and many of the genetic regulators of lifespan have been linked to lipid metabolism (Mutlu et al., 2021). We sought to determine lipid abundance changes in the same cortical tissue samples used for transcriptomic analysis and identify if lipid metabolism associated alternative splicing events were enriched in response to CR. We used shotgun LC-MS/MS lipidomics, identifying 721 total lipid species combined between positive and negative mode ionization. Of these, 549 lipid species passed filtering thresholds and were quantified. Focusing on the response to CR, we found 145, 134, and 137 statistically significant lipid species at 10 months, 20 months, and 30 months of age, respectively (**Fig.5A; Table S7**). Comparing these lipids collectively, we found substantial overlap among these lipids, with 24% shared across the diet response at all three ages, and 55% shared by at least two age groups (**Fig.5B**); this suggests a consistent CR effect independent of age, similar to the transcript DEG results. Using PCA, we observe a similar separation of samples by diet on PC 1, also like the transcriptome (**Fig.5C**). However, here the effect of age is also visible in the lipid PCA, largely confined to PC3. This is potentially the result of an age effect on lipid composition that is more overt than the larger feature space transcriptional dataset.

**Fig. 5.**
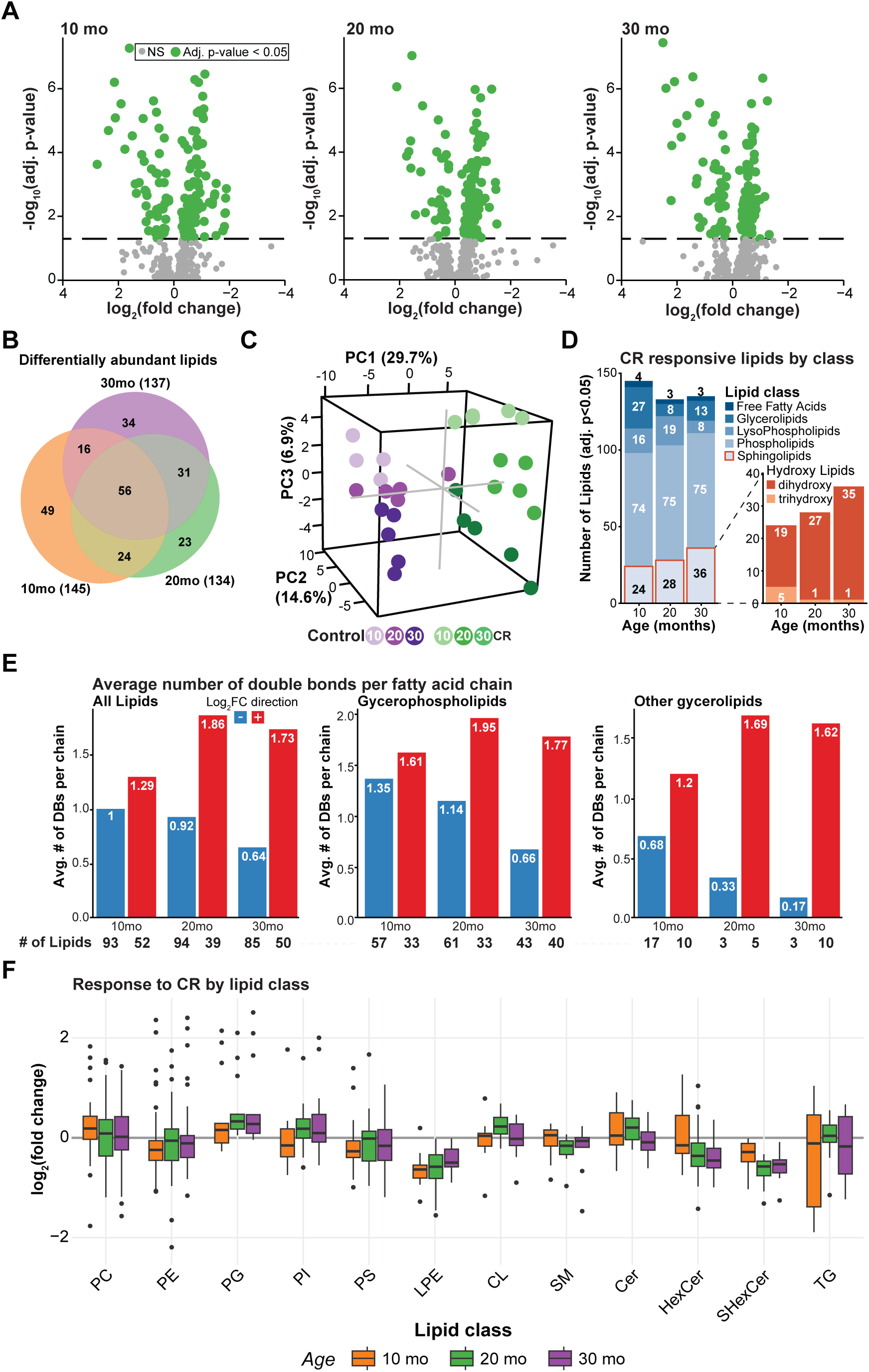
Caloric restriction alters lipid composition in the brain in an age-dependent manner. (**A**) Volcano plots depicting differential lipid abundance in response to diet at 10, 20, and 30 months; lipids significant at adjusted p < 0.05 are shown in green. (**B**) Euler diagram of the differentially abundant lipids (adjusted p < 0.05) shared among the three ages. (**C**) Principal component analysis of the lipidome (PC1, 29.7%; PC2, 14.6%; PC3, 6.9%), with samples colored by diet and age (control and CR at 10, 20, and 30 months). (**D**) Number of CR-responsive differentially abundant lipids (adjusted p < 0.05) at each age, colored by lipid class (free fatty acids, glycerolipids, lysophospholipids, phospholipids, sphingolipids). Inset: among the significant hydroxylated sphingolipids, the number of dihydroxy-versus trihydroxy-sphingoid-base species per age. (**E**) Average number of acyl-chain double bonds among CR-decreased (blue; negative log_2_ fold change) and CR-increased (red; positive log_2_ fold change) lipids, by age, for all lipids and for the glycerophospholipid and glycerolipid subsets; the number of lipids per group is listed below each panel. (**F**) Distribution of the log_2_ fold change (restricted versus control) for each lipid class, by age (10, 20, and 30 months).

The bulk of CR-responsive lipids were phospholipids, comprising over half of all significant changing lipid species at each age (**Fig.5D**), a proportion that was consistent regardless of age. In contrast, the proportion of CR responsive lipids identified as sphingolipids increased with age, from 17% of significantly changing lipids at 10 months to 21% at 20 months to 27% at 30 months of age. All significant CR responsive sphingolipids were ceramide species. Importantly, CR opposed the age-related increase in brain resident ceramide species and all 36 ceramide species detected were downregulated due to CR at 30 months. In circulation, many reports point to associations between increased ceramide concentrations and age-related diseases (Chan et al., 2018; Smith et al., 2022). Our data is consistent with these trends that link lower ceramide accumulation to healthier outcomes; however, it is not clear what reductions in ceramide levels within a tissue might indicate. Individual cell types within the brain have substantially different contributions and sensitivities to ceramide levels (McInnis et al., 2024), so more work will be necessary to understand the implications of these observations.

We next examined fatty acid chain unsaturation levels. Fatty acid double bonds can impact cellular features such as membrane fluidity while also functioning as a potential site of damage from free radicals and lipid peroxidation (Riahi et al., 2010). Across all lipids there is a strong separation of average chain unsaturation level by lipid fold change in response to diet. This effect gets stronger with age: at 10 months, the average number of double bonds across all chains is close to one, and there is a 0.29 average double bond difference between downregulated and upregulated lipid species (**Fig.5E**); at 20 months, this has widened to a 0.94 difference, with the average number of double bonds among upregulated lipid species at 1.86 compared to 0.92 for downregulated lipids; at 30 months of age, the gap is 1.09 double bonds on average. A similar trend can be observed if we examine subgroup classes of lipids, such as the glycerophospholipids and other glycerolipids. Overall, this indicates an increase in the unsaturation index across the identified lipids, which is consistent with previous reports (Laganiere and Yu, 1987).

To uncover CR-responsive trends across lipid classes, we performed Lipid Set Enrichment Analysis (LSEA). This process ranks all lipid species by diet-response and then examines enrichment of particular categories of lipids towards the top or bottom of the list. The resulting analysis detected subclasses of glycerosphingolipids, specifically hexosylceramides (HexCer) and sulfonated hexosylceramides (SHexCer), and lysophosphatidylethanolamines (LPE) as enriched among downregulated lipid species (**Fig.5F**). Hexosylceramides (cerebrosides) and phosphatidylethanolamines are major components of myelin, which is critical for proper nervous system function. Remodeling of myelin lipid composition may contribute to established benefits of CR on brain function and suppression of neurodegenerative disease development (Babygirija et al., 2025; de Oliveira et al., 2022; Halagappa et al., 2007).

### Regularized canonical correlation analysis identifies shared molecular signatures

To identify the degree of correlation across data types, we integrated the transcript expression, splicing, and lipid composition datasets using canonical correlation analysis. We first used regularized canonical correlation analysis (rCCA) to examine the correspondence between each datatype in a pairwise manner (**Fig.6A&B**). The process for rCCA as performed here relies on principle component analysis for each dataset individually, which are then blended pairwise in a weighted manner such that the weights are determined by what maximizes the correlation between each data type, thereby identifying a shared axis of variation. Post-hoc regression of the resulting scores against diet or age sheds light on the phenotypic driver(s) of this shared axis. Examining the individual sample scores for each aspect in box plots, separating samples by diet group, we can observe a clear diet-driven signal in the first shared component between the transcript and lipid datasets (**Fig.6A**, *middle*). Component 2 and 3 showed almost no separation by diet among the sample scores, suggesting that the diet effect is almost entirely captured in the first shared axis (**Table S8**). This general trend is also true of the transcript-splicing (**Fig.6B**) and lipid-splicing rCCA comparisons. We can then draw inferences about the molecules driving this shared axis by ranking the members of each dataset by their respective correlation with the shared axis (**Table S9**). For the transcript-lipid rCCA, we performed GSEA Reactome pathway enrichment based on the list of transcripts ranked by correlation magnitude (from strongest correlation, regardless of sign, to weakest correlation) (**Fig.6A**, *left*). We found that the strongest correlated factors were enriched for many pathway categories that classically respond to CR: transcription and translation, metabolism, and signal transduction and growth pathways (**Fig.6A**). This shared axis therefore suggests a possible mechanistic connection between lipid composition and gene expression regulation that was not apparent from transcript differential expression analysis alone. To examine the lipids in this comparison in a similar fashion, we performed LSEA on the corresponding list of lipids ranked based on correlation with the shared axis, finding that the strongest correlated lipid species were enriched for monounsaturated fatty acyl chains and di-hydroxy sphingoid base (**Fig.6A**, *right*). Sphingolipids, ceramides, phospholipids and glycerolipids showed substantial enrichment as well, but did not reach statistical significance. The mechanistic connection between lipid composition and gene expression regulation in response to CR may therefore involve monounsaturated fatty acids and/or di-hydroxy sphingolipids.

**Fig. 6.**
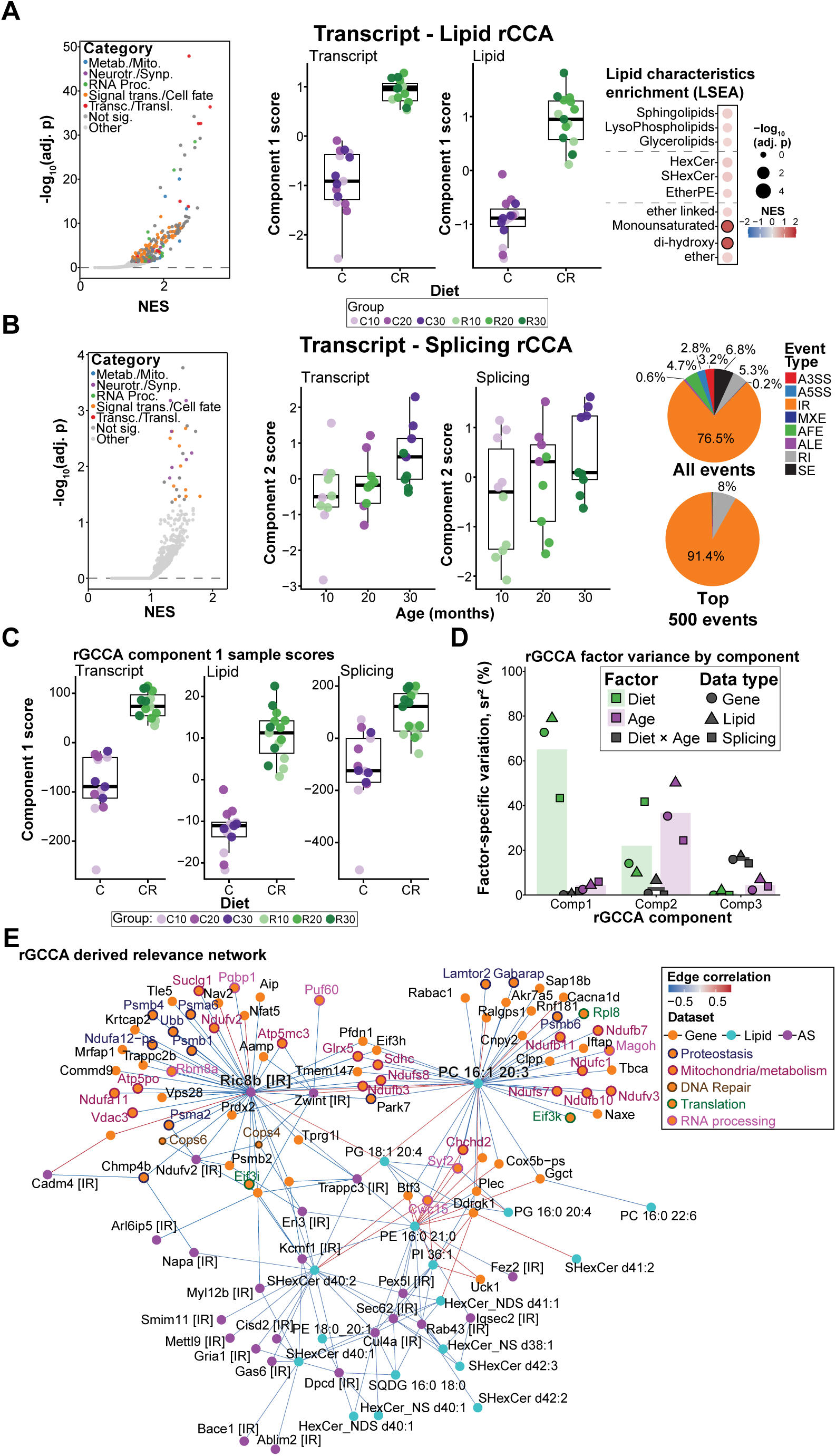
Canonical correlation analyses emphasizes shared variation in response to diet and impact of intron retention. (**A**) Transcript-lipid pairwise rCCA, component 1. *Left*: volcano plot depicting gene-set enrichment (GSEA) of the transcript component-1 ranking; pathways are colored by functional category (Metabolism/Mitochondria; Neurotransmitter/Synaptic; RNA processing; Signal transduction/Cell fate; Transcription/Translation; Other), non-significant pathways in grey. *Middle*: per-animal component-1 scores for the transcript and lipid datatypes, separated by diet. *Right*: lipid-characteristics enrichment (LSEA) of the component-1 lipid ranking. (**B**) Transcript-splicing pairwise rCCA, component 2. *Left*: volcano plot depicting GSEA of the transcript component-2 ranking (axes and category coloring as in **A**). *Middle*: per-animal component-2 scores for the transcript and splicing datatypes, separated by age (10, 20, 30 months). *Right*: SpliceWiz event-type composition of the splicing component-2 ranking, for all events and for the 500 most strongly ("top") correlated events; slices are colored by event type (IR, RI, SE, AFE, ALE, A5SS, A3SS, MXE). (**C**) Per-animal rGCCA component 1 scores for each of the three datatypes, separated by diet. (**D**) Squared semipartial correlation (sr²) is shown as the percentage of total component-score variance uniquely associated with the factors Diet, Age, or the Diet × Age interaction. Points show the datatype-specific values for transcript, lipid, and splicing datatypes, and bars show their equal-weight descriptive mean. Diet is shown in green, age in purple, and diet × age in grey; the interaction is displayed as the three datatype-specific points with a bar indicating the mean. (**E**) Relevance network derived from the rGCCA analysis. Nodes are the top component-1 (diet) features of each layer - genes, lipids, and alternative-splicing (AS) events - colored by data layer and sized by their absolute correlation with the component-1 variate; gene nodes are additionally labeled by functional category (Mitochondria/metabolism, Proteostasis, Translation, DNA repair, RNA processing). Edges connect features of different layers based on pairwise correlation and are colored by the sign and strength of the cross-layer correlation between them (red, positive; blue, negative).

Although age as a variable was generally less resolved onto one axis and thus less visible in these analyses, the shared component 2 of the transcript-splicing rCCA provides evidence of an age or age-diet combined effect (**Fig.6B**, *middle*). The scores for the transcript aspect increase slightly from 10 months to 20 months, and then more dramatically at 30 months of age. A less defined pathway response was observed using the list of transcripts ranked by correlation with component 2, with fewer pathways representing our manually-defined shared pathway categories (**Fig.6B**, *left*; **Table S10**); this is consistent with our transcript DEG analysis, where a strong age signature was not observable. For the alternative splicing aspect of component 2, where the individual sample scores indicate a diet-by-age effect, the overall list of alternative splicing events was dominated by intron retention events (comprising 77% of the input), but are especially enriched (91%) in the events most strongly correlated with the shared axis (**Fig.6B**, *right*). This argues that intron retention events are substantially related to both diet AND age in this context, something that was not visible in our conventional differential splicing analysis (**Fig.3**).

We performed regularized generalized canonical correlation analysis (rGCCA) to examine shared correlation across all three datasets. As its name indicates, rGCCA is a generalization of rCCA to three or more datatypes. Examining the sample scores for component 1 across all three datatypes, we observed a substantial difference by diet in all three, similar to the paired rCCA comparisons, although the splicing event data showed the smallest separation (**Fig.6C**). We used squared semipartial correlation, which quantifies the proportion of variance uniquely associated with a set of predictors, to examine the amount of variance across the data that was confined to each component and data type (**Fig.6D**). Component 1 is strongly associated with diet across all three data types, with diet explaining roughly 65% of the shared variance. Approximately 40% of shared variance across component 2 is associated with age, matching results from the pairwise rCCA analyses (**Fig.S2**). To identify potential molecular drivers of the CR response, we then ranked all contributing features to the rGCCA by Pearson correlation with component 1. Selecting the top molecules from each data type, proportionally to each input dataset size, we constructed a relevance network (**Fig.6E**). Nodes represented the top molecular features correlating with the rGCCA component 1, while edges represented the Pearson correlation between each molecule pair; edges were not calculated between nodes derived from the same data type. Examining the network structure, two semi-defined clusters are apparent: a group of transcripts centered around an intron-retention event in the gene Ric8b, with which the transcripts are mostly negatively correlated, and a similarly arranged cluster centered around a phosphatidylcholine with two unsaturated fatty acyl chains. Ric8b is a guanine nucleotide exchange factor involved in G protein-coupled receptor signaling, while phosphatidylcholine is likely to be a membrane component. In both cases, the central position of each factor within the network suggests a fundamental role in the CR response. The bottom half of the network is comprised of a less strongly correlated group of IR events and a variety of lipid species, especially ceramides. Overall, the network contains a large number of factors associated with the mitochondrial electron transport chain (especially complex I) and proteostasis. The lipid signature is dominated by ceramides, consistent with our differential abundance analysis, while the alternative splicing events included in the network are exclusively IR events, a feature that was less apparent in the differential splicing analysis.

## Discussion

In line with previous reports (Heintz et al., 2017; Rhoads et al., 2018; Seo et al., 2016), the data presented here provides evidence that altered RNA processing is a central contributor to the mechanisms of CR. We found that the gene expression response to CR is relatively consistent regardless of life-stage of the animals, including general downregulation of the machinery responsible for most of the aspects of RNA processing. In contrast to the robust diet response, we found relatively few differentially expressed transcripts in response to age. This is likely due to a combination of factors including gene expression levels being relatively stable in response to age when examined via bulk RNA sequencing, small effect size changes in response to age, increased group variance with age (Salimi et al., 2026), and the cell-type composition complexity of brain tissue (Yao et al., 2023). Pathway analysis revealed upregulation of neuronal function and synaptic structure-related pathways. Pathways normally associated with upregulation in response to CR in other tissues were downregulated here, including transcriptional processing and metabolism/mitochondria pathways. Although an inversion of what has been reported for the CR response in other tissues, the direction of effect here is the opposite of what has been noted in the brain for the effect of age, suggesting CR is contributing to the maintenance of a more youthful transcriptional program (Souder et al., 2025). It would be of considerable interest to identify the dynamics of this transcriptional response across all contributing cell types, accounting for cell-type changes with age, at full transcriptome depth and thereby enabling a more complete picture of the gene expression regulation.

Corresponding to the general downregulation of the transcriptome, we found that much of the machinery responsible for expression and processing of mRNA transcripts was sharply downregulated at the transcript level, including transcription machinery, the spliceosome, export factors, and protein coding ribosomal transcripts. In part, this homeostatic shift could be a response to nutrient limitations reducing transcript, and thus protein, production which is an energetically demanding processes (Qiao et al., 2024). In parallel, the primary energy production pathways are downregulated at the transcriptional level. Interestingly, when looking at the properties of the individual transcripts being produced, we found that there was a bias in favor of upregulation of the longest transcripts that would be predicted to be the most energetically expensive to produce. This is counter to the transcriptional shortening that is observed with age (Stoeger et al., 2022), and overall points to a distinct and programmatic remodeling of the transcriptome in response to CR rather than simply a global reduction of transcription and protein production. Here also it will be important for future research to define which cell types are most responsible for these effects.

We observed distinct changes to lipid composition in response to CR, notably decreases in ceramide levels and remodeling of the distribution and number of double bonds among fatty acid chains. Ceramide species have been associated with aging and circulating ceramides have been explored as biomarkers of age-associated diseases such as cardiovascular disease (McGurk et al., 2021). Our data suggest that reduced ceramide levels are a feature of CR, consistent with other reports. A surprising observation, however, was the increase in unsaturation numbers among upregulated lipid species, given the potential for an increased risk of lipid peroxidation with increased prevalence of double bonds; increased lipid peroxidation is a noted feature of aging (Praticò, 2002). However, as our dataset is largely comprised of phospholipids that are primarily components of lipid membranes, it’s possible that increased unsaturation is necessary to counteract membrane fluidity declines due to age. Further work will be necessary to understand both the drivers and the consequences of these changes with regards to enhanced longevity.

The most notable signature observed in our analysis of alternative splicing is increased intron retention. In the 30-month old animals, IR events comprise a large fraction of the statistically significant events identified in response to diet (**Fig.3B**); however, our canonical correlation integration strategy uncovered a much more widespread IR signature, as IR events dominate the rCCA/rGCCA shared axes and the resulting relevance network (**Fig.6**). Interestingly, the vast majority of these events are increased retention in response to diet. This is consistent with the global downregulation of the splicing and transcriptional processing machinery leading to less splicing in response to CR; however, increased intron retention is associated with progressive aging and age-associated diseases such as Alzheimer’s (Adusumalli et al., 2019), so it is unclear how this increase might be linked to beneficial aging effects. Most of the relevant events are predicted to be candidates for nonsense mediated decay, suggesting this may be a mechanism of expression downregulation and pathological aging may be associated with different intron retention events. However, it will be important to establish the specific consequences, cell-type origins, and relevance for the healthspan/lifespan benefits for individual events in future work.

The canonical correlation analyses revealed key integrated aspects of the CR response. The relevance network represents the factors most correlated with the canonical axis that aligns with the diet response, and includes transcripts associated with pathways known to be harnessed by CR – metabolism, proteostasis, DNA repair, translation, and RNA processing. These factors largely form two subnetworks centered around an IR event in the Ric8b gene and a phosphatidylcholine species. Additionally, the bottom 1/3 of the network highlights a cluster of ceramides correlated with a variety of IR splicing events, suggesting a possible relationship between ceramide levels and splicing. Both regulation of splicing by ceramides (Chalfant et al., 2002; Massiello et al., 2004) and regulation of ceramide levels by splicing (Pani et al., 2021) have been described, indicating a potential reciprocal relationship. These features highlight aspects of the CR response that have yet to be mechanistically investigated and are thus promising targets for uncovering novel mediators of the beneficial effects of CR.

Overall, our data suggests that changes to splicing are widespread in response to CR, affecting many aspects of the cellular response, and further investigation of the regulation of alternative splicing in response to delayed aging interventions has a high likelihood of uncovering mechanistic drivers.

### Limitations

There are several aspects of this study that limit the generalizability of the conclusions and warrant future investigation. As a result of working from banked tissue from a previous study, our analyses are from male mice only; it will be important to perform similar analyses in female mice in the future. The study design is cross-sectional, so we are limited in our ability to define full trajectories for the observed changes. Like with many conventional CR implementations, there is a substantial fasting period that may drive many observed changes, and mice are singly-housed for the purpose of careful tracking of food intake, which may impact social modifiers of mouse physiology. Finally, although we perform cell-type deconvolution to partially address this, many of the observed changes are likely cell-type specific but we are not able to granularly assign them because our data is derived from bulk RNAseq. Future improvements in single-cell sequencing may enable deep alternative splicing analysis in the future to address this.

## Methods

### Animal Model and Study Design

This study used banked flash-frozen cortex from a cohort of male B6C3F1 hybrid mice (Miller et al., 2017; Souder et al., 2025). The original study was approved by the Institutional Animal Care and Use Committee at the University of Wisconsin-Madison. Briefly, mice were housed under pathogen-free conditions. At 2 months of age, mice were randomized into control or restricted diet groups, fed 87 kcal/wk (95% of *ad libitum* intake) (Bio-Serv diet #F05312), or 73 kcal/wk (Bio-Serv diet #F05314), respectively. The restricted diet is a 20% reduction in caloric intake from *ad libitum* levels and 16% reduction from controls. Mice were euthanized by cervical dislocation at 10, 20, or 30 months of age. For the purposes of the original study, brain tissue was isolated, embedded in OCT (Fisher Scientific), and frozen in liquid nitrogen and stored at -80 °C until further processing.

### Tissue Collection and RNA extraction

For the present study, 50-100 mg of frozen cortical tissue was punched out of the OCT embedded brain tissue using a 1-mm biopsy punch (Integra Miltex). The resulting tissue punch was ground in 0.8 mL of Trizol (ThermoFisher, #15596026) using a Polytron homogenizer. To induce phase separation, 0.1X volumes of 1-chloro-2-bromopropane was added to the Trizol-tissue homogenate and spun at 12,000 x *g* at 4 °C for 15 minutes. The aqueous phase containing RNA was moved to a new tube and RNA was extracted using a Direct-zol RNA Miniprep (Zymo Research, #R2052) according to the manufacturer’s instructions. Total RNA integrity was measured on an RNA Pico Chip using a Bioanalyzer 2100. 1 μg of total RNA was used as the input for library construction.

### Transcriptomics

Poly-A transcripts were enriched from total RNA using the NEBNext Poly(A) mRNA Magnetic Isolation Module (NEB, #E7490L). Libraries were then constructed using NEBNext Ultra RNA Library Prep Kit for Illumina (NEB, #E7530L) with dual-index barcodes (NEB, #E6330S). The libraries were sequenced 2×150 bp to a depth of approximately 100 million reads per sample on an Illumina NovaSeq 6000. Raw reads in FASTQ format were aligned to the mouse reference genome (ENSEMBL GRCm39) using Rsubread. Reads were not trimmed (Williams et al., 2016). Gene-level read counts were obtained with Rsubread::featureCounts against the Mus_musculus.GRCm39.113 GTF annotation.

Differential gene expression was performed with edgeR in combination with limma-voom (Ritchie et al., 2015). Raw read counts were filtered and TMM-normalized. Voom-transformed log-CPM values were then fit against a no-intercept design matrix including the six experimental groups (C10mo, R10mo, C20mo, R20mo, C30mo, R30mo). Contrast matrices were built on age (C20-C10, C30-C20, C30-C10, R20-R10, R30-R20, R30-R10) and diet (R10-C10, R20-C20, R30-C30). Empirical Bayes shrinkage was applied and full ranked gene lists were extracted; transcripts were considered differentially expressed if the adjusted p-value was less than or equal to 0.05, with additional criteria of absolute magnitude of log_2_ fold-change greater than 0.75 used where noted.

### Pathway analysis

Gene Set Enrichment Analysis (GSEA) was performed using the WebGestalt web tool (Liao et al., 2019) against the Reactome mouse pathway database. For all cases where pathway analysis was used, significantly enriched Reactome pathways were grouped into a fixed set of functional categories defined for this study: Transcription and Translation; Metabolism and Mitochondria; RNA Processing; Neurotransmission and Synaptic Signaling; and Signal Transduction and Cell-Fate Regulation. Pathways that did not clearly belong to one of these categories were assigned to Other. Every significant pathway was categorized manually by review of its biological function - its Reactome annotation and the process it represents. Pathways with plausible membership in more than one category were flagged, reviewed, and assigned to their predominant function.

### Transcript network analysis

Weighted Gene Co-expression Network Analysis (WGCNA) (Langfelder and Horvath, 2008) was applied to a TMM-normalized log-CPM expression matrix consisting of 21,729 input genes. A signed adjacency matrix was constructed using a soft-thresholding power chosen as the smallest value satisfying a scale-free topology fit of R^2 ≥ 0.8. Modules were identified using signed network and TOM types, Pearson correlation, minModuleSize = 30, mergeCutHeight = 0.25, and maxBlockSize = 20,000. A random seed was not set for module detection, so block-boundary module assignments are not bit-reproducible across runs. No explicit batch-correction step was applied prior to network construction. Module eigengenes were correlated against the following traits: diet (CR vs. control), age (in months), and the interaction of those two terms. Significance of correlations was determined by FDR < 0.05.

### Module-Pathway over-representation analysis

For each non-gray WGCNA module, the member gene list was tested for Reactome pathway enrichment using clusterProfiler (Yu et al., 2012). Ensembl gene identifiers were mapped to Entrez identifiers via org.Mm.eg.db and deduplicated. Enrichment was run with organism = “mouse”, gene-set size restricted to between 15 and 500, and background gene set included all WGCNA-input genes that successfully mapped to Entrez identifiers. Modules with fewer than 10 Entrez-mapped genes were skipped. Pathways with adjusted p < 0.05 were retained.

### Curated genesets

Gene membership in the spliceosome category was manually curated, guided by the literature and existing databases (Cvitkovic and Jurica, 2013). Human gene symbols were converted to mouse orthologs using the Mouse Genome Informatics HOM_MouseHumanSequence homology table, restricted to one-to-one ortholog groups, and combined with an independent biomaRt-based conversion of the same human source sets before deduplication. The resulting mouse-symbol list was divided into 158 core spliceosome genes and 413 regulatory factors on the basis of (Papasaikas et al., 2015) – 510 of these 571 were observed in our transcriptomics data. Membership in the ribosome category was based on HGNC gene symbol conventions (RPL for large subunit, RPS for small subunit). Membership in the transcription and export categories was based on literature (Blombach et al., 2016; Katahira, 2012).

### Cell-type Deconvolution

Bulk RNA-seq expression was deconvolved into estimated brain cell-type signatures using BrainInABlender with species = "mouse" (Hagenauer et al., 2018). Raw counts were TMM normalized using edgeR; Ensembl identifiers were mapped to mouse gene symbols via org.Mm.eg.db. Average primary cell-type indices were extracted per sample and visualized as pheatmap heatmaps (blue-white-red colormap, row-clustered, columns ordered by experimental group). Inter-sample variance of cell-type marker z-scores was quantified across age and diet groups.

### Alternative Splicing Events

Alternative splicing events were detected from the aligned RNAseq reads using the R package *SpliceWiz* (Wong et al., 2024). Two control samples consistently failed SpliceWiz processing and were excluded from downstream analysis, leaving 28 of 30 samples (with no fewer than 4 samples per group). Analysis was performed for diet contrasts (10-month R vs. C: 5 control and 5 restricted samples; 20-month R vs. C: 4 control and 5 restricted samples; 30-month R vs. C: 4 control and 5 restricted samples). Default event-quality filters were applied, and Percent Spliced In (PSI) values were computed as Included / (Included + Excluded) from the assay matrices. Events with Benjamini-Hochberg FDR < 0.05 were considered differentially spliced. Event-type categories considered were: skipped exon (SE), intron retention (IR), retained intron (RI), alternative 3’ splice site (A3SS), alternative 5’ splice site (A5SS), alternative first exon (AFE), mutually exclusive exon (MXE) and alternative last exon (ALE). To minimize confusion, RI events, which substantially overlap with IR events but are calculated differently than the remaining event types, are only included in unsupervised analyses but ignored for downstream analyses. For examination of intron retention, we focus on IR events.

### Splicing WGCNA

Per-event PSI matrices were obtained from SpliceWiz across seven alternative-splicing event types (A3SS, A5SS, AFE, ALE, IR, MXE, and SE). For each event type, only events quantified across all samples were included. Events were filtered according to the following criteria: events containing NAs were removed, and for feasibility of calculating the WGCNA matrix, the remainder were ranked based on row-wise median absolute deviation to focus on the most variable 30,000 events. These events were logit-(log-odds) transformed, with a single global offset (half the smallest interior PSI value observed across the matrix) added to or subtracted from exact values of 0 and 1, respectively, prior to transformation.

A signed co-expression network was built using Pearson correlation and a signed topological overlap matrix in a single block. The soft-thresholding power was chosen as 8. Modules were called with a minimum module size of 30 and were merged at an eigengene-dissimilarity height of 0.25. Module eigengenes were correlated (Pearson) with six traits: diet (restricted vs. control), continuous age, indicator variables for each age group (10, 20, and 30 months), and a diet-by-age interaction. Student’s t-test derived p-values were adjusted across all module-trait pairs by the Benjamini-Hochberg procedure, and associations with FDR < 0.05 were considered significant. For every module and event-type combination, enrichment of that event type within the module relative to the rest of the network was tested by Fisher’s exact test, adjusted with Benjamini-Hochberg FDR across all combinations.

### Lipidomics sample preparation

gDNA was precipitated from the organic phase leftover from Trizol-based RNA extraction (see *Tissue Collection and RNA extraction*, above) with the addition of 0.3X volumes of 100% ethanol followed by centrifugation at 5,000 x *g* at 4 °C for 5 minutes. The supernatant was moved to a new tube and proteins precipitated by the addition of 1.5X volumes of 100% 2-propanol followed by centrifugation at 12,000 x *g* at 4 °C for 10 minutes. The supernatant containing lipids was moved to a new tube, dried under nitrogen to reduce volume to ≤ 500 μL, and lipids subsequently isolated via the Folch method using chloroform:methanol (Folch et al., 1957). The remaining organic fraction after extraction was dried under liquid nitrogen and resuspended in 300 μL of LC-grade isopropanol in preparation for LC-MS.

### Lipidomics LC-MS

Lipidomics analyses were performed by the University of Wisconsin Biotechnology Center Mass Spectrometry Facility. Lipid extracts were separated on an Agilent InfinityLab Poroshell 120 EC-C18 1.9 µm 2.1 x 50 mm column maintained at 50 °C connected to an Agilent HiP 1290 Multisampler, Agilent 1290 Infinity II binary pump, and column compartment connected to an Agilent 6546 Accurate Mass Q-TOF dual ESI mass spectrometer. For positive mode, the source gas temperature was set to 250 °C, with a gas flow of 12 L/min and a nebulizer pressure of 35 psig. VCap voltage was set at 4000 V, fragmentor at 145 V, skimmer at 45 V and Octopole RF peak at 750 V. For negative mode, the source gas temperature was set to 350 °C, with a drying gas flow of 12 L/min and a nebulizer pressure of 25 psig. VCap voltage was set at 5000 V, fragmentor at 200 V, skimmer at 45 V and Octopole RF peak at 750 V. Reference masses in positive mode (m/z 121.0509 and 922.0098) and negative mode (m/z 1033.988, 966.0007, and 112.9856) were delivered to the second emitter in the dual ESI source by isocratic pump at 15uL/min.

Samples were analyzed in a randomized order in both positive and negative ionization modes in separate experiments acquiring with the scan range m/z 100 – 1500. Mobile phase A consisted of ACN:H2O (60:40 v/v) containing 10 mM ammonium formate and 0.1% formic acid, and mobile phase B consisted of IPA:ACN:H2O (90:9:1 v/v) containing 10 mM ammonium formate and 0.1% formic acid. The chromatography gradient for both positive and negative modes started at 15% mobile phase B then increased to 30% B over 2.4 min, it then increased to 48% B from 2.4 – 3.0 min, then increased to 82% B from 3 – 13.2 min, then increased to 99% B from 13.2 – 13.8 min where it was held until 15.4 min and then returned to the initial conditions and equilibrated for 4 min. Flow was 0.5 mL/min throughout, injection volumes were 1µL for positive and 5 µL for negative mode. Tandem mass spectrometry was conducted using the same LC gradient at collision energy of 25 V.

QC samples and blanks were injected throughout the sample queue and ensured the reliability of acquired lipidomics data. Results from LC-MS experiments were collected using Agilent Mass Hunter (MH) Workstation Data Acquisition. Briefly, a pooled lipid extract comprised of an aliquot from each sample was analyzed in MS/MS mode. From this data, a lipid library was created using Lipid Annotator (Agilent Technologies, Inc) for positive and negative ion data, incorporating lipid identity (with either enumerated acyl chain composition or a sum composition), m/z value, and retention time. Data for all individual samples was collected in MS mode. Profinder (Agilent Technologies, Inc.) was used to perform retention time alignment between samples and to extract peak areas from each sample, for each lipid present in the library generated by Lipid Annotator.

### Lipidomics data analysis

Raw peak-area tables were curated to disambiguate chromatographic duplicate peaks, remove low-quality peaks flagged during manual chromatogram review, and correct known impossible values. Lipid names were tagged with their ionization mode to keep mode-distinct peaks separate during pooled analysis. Per-row missingness was computed across the 30 samples and lipids with more than 20% missing values were dropped. Remaining missing values were imputed at half the lipid row’s minimum observed value (half-minimum imputation) (Frölich et al., 2024). Imputed abundances were log_2_-transformed and per-sample median-centered, and positive- and negative-mode matrices were then combined to form the combined lipid abundance matrix.

For each lipid, a two-way analysis of variance (abundance ∼ group * age) was fit and followed by Tukey’s HSD post-hoc comparison. Log_2_ fold changes were computed from per-group sample means on the normalized abundance scale. Lipids with Tukey adjusted p-value < 0.05 in any R-versus-C contrast were considered differentially abundant.

Each lipid species name was parsed to assign a headgroup class label (glycerophospholipid, lysophospholipid, sphingolipid, fatty acid, and others); within the sphingolipid group, ceramide species were further partitioned by sphingoid-base hydroxylation state (dihydroxy versus trihydroxy) using the d/t prefix conventions in the ProFinder lipid identifiers. For acyl-chain saturation, the C:D suffix(es) of each significant lipid (number of carbons and number of double bonds) were parsed.

Lipid set enrichment analysis (LSEA) was performed using the R package lipidr (Mohamed et al., 2020). Lipids were grouped into characteristic sets defined as in the study’s lipidomics pipeline: major classes (Phospholipids, LysoPhospholipids, FattyAcyls, Glycerolipids, Sphingolipids) and lipid-composition sets (saturation class and fatty-acid content). For each CR vs control contrast, all detected lipid species were ranked by their log_2_ fold change (positive- and negative-ionization-mode measurements were kept as distinct features and ranked together in a single list) and lipid set enrichment was computed with fgsea::fgseaMultilevel on the fold change ranked list, yielding a normalized enrichment score (NES) and a Benjamini-Hochberg false-discovery rate adjusted p-value for each set.

### Multiomic Integration

We performed regularized canonical correlation analysis (rCCA) using the mixOmics R package. This analysis was performed pairwise between each combination of data types (transcript – lipid, transcript – splicing, lipid – splicing). Because every layer holds far more features than samples, the layers cannot be correlated feature-by-feature in a stable way, so each layer was first summarized on its own. Following centering and scaling, principal component analysis was performed. This process replaced the individual molecular features of a layer with eight principal components. For the splicing data, percent-spliced-in values were transformed using a symmetric epsilon and then logit-transformed, as described in the splicing WGCNA section above. rCCA was then performed on the resulting PCs, retaining three shared components per rCCA in the output.

Each pair of layers was analyzed by regularized canonical correlation analysis (mixOmics::rcc, method = "shrinkage", ncomp = 3). For a pair of layers, the method searches each layer’s eight summary scores for the weighted combination that, matched against the corresponding combination in the other layer, lines up as closely as possible across animals, giving each animal one combined score per layer that captures the pattern the two layers most share (the first three such combinations were kept). Association with individual molecules was determined by how strongly each feature’s own measured values correlated with the shared axis score across animals (signed Pearson correlation).

To describe all three datasets together, the PCA-derived layers were fit jointly by regularized generalized canonical correlation analysis (mixOmics::wrapper.rgcca), using the 28 animals shared across all three datasets. For each of three retained components, every layer receives one combined score per animal, but, unlike the pairwise method, each score is chosen to agree with both of the other two layers at once (Horst, 1961). To quantify factor-specific variation in the rGCCA components, block-specific component scores for gene expression, lipidomics, and alternative splicing were treated as per-animal response variables. Modeling canonical variate scores as outcomes of experimental factors has precedent in CCA-based analyses (Krimmel et al., 2022). For each data type block and each rGCCA component, a two-way factorial linear model was fit including diet (Control or CR), age (10, 20, or 30 months), and the diet × age interaction. Type III sums of squares were calculated using sum-to-zero contrasts, allowing each model term to be evaluated while accounting for the remaining terms (Hector et al., 2010). Factor-specific (factor meaning sample variable – diet, age, or interaction terms) variation was quantified using squared semipartial correlation (sr²), calculated as the Type III sum of squares for each term divided by the total corrected sum of squares of the component score. Semipartial R² quantifies the proportion of outcome variance uniquely associated with a predictor or set of predictors and can be calculated for multilevel factors and interaction terms treated as predictor sets (Stoffel et al., 2021). Gene, Lipid, and PSI sr² values were plotted individually, with their equal-weight mean shown as a descriptive summary across the three data type blocks. As in the pairwise rCCA analysis, we then ranked all features by correlation with its layer’s combined score for the first component, retaining a scaled number of features for each datatype based on total input (log-proportionally: 107 transcripts, 67 lipids, and 126 splicing events); these features were then used as nodes to construct a relevance network. Edges were drawn between nodes of different data types (never within a data type) from the direct correlation of the two features, retaining the strongest cross-layer correlations for each layer-pair, and the network was laid out with a Fruchterman-Reingold algorithm, which places strongly correlated features near one another.

### Data Analysis Pipelines

All R analyses were performed in R 4.5.3 on Windows 10 x64 (build 19045). Key R packages used in this study: Rsubread (alignment and counting); edgeR and limma (gene-level differential expression and voom transformation); SpliceWiz (alternative splicing detection and event-level testing); WGCNA (gene co-expression network analysis); splicejam (sashimi-style visualization of alternative splicing events); EnhancedVolcano (volcano plots); eulerr (Euler-style Venn diagrams); ComplexHeatmap and pheatmap (heatmaps); rgl (three-dimensional scatter plots); mixOmics (block PCA and regularized canonical correlation analysis); ReactomePA and clusterProfiler (Reactome over-representation analysis on WGCNA modules); BrainInABlender (cell-type deconvolution). External web tools: the WebGestalt server (gene set enrichment analysis on the gene expression layer; over-representation analysis on the MixOmics co-varying-block gene members).

### Statistical Analysis

Unless otherwise stated, multiple hypothesis testing used Benjamini-Hochberg false discovery rate control, and the significance threshold was FDR < 0.05. For per-lipid two-way ANOVA, the Tukey HSD adjusted p-value was used as the significance criterion. For rCCA component-by-trait labelling, an uncorrected alpha = 0.05 cutoff was used on the diet/age/interaction linear-model coefficients.

## Author Contributions

SAT and TWR designed the experiments. The original mouse study from which the samples used herein were derived was designed by RMA. JPC and TWR collected and processed samples. SAT, AJE, AS, JPC, and TWR performed data analyses. TWR supervised the project. SAT and TWR wrote the manuscript in consultation with all authors. All authors contributed to editing and revising of the manuscript.

## Acknowledgements

We thank Dr. Samantha Waters for assistance with sample acquisition. We acknowledge funding from the NIH/NIA (R01AG037000 to RMA) and the University of Wisconsin Comprehensive Diabetes Center pilot award (UWCDC-CSPA-20-4 to TWR), as well as the Office of the Vice Chancellor for Research at UW-Madison (to TWR). RMA and TWR are members of the Wisconsin Nathan Shock Center of Excellence in the Basic Biology of Aging, NIH/NIA P30AG092586. We thank the UW-Madison Biotechnology Center Gene Expression Center and DNA Sequencing Facility for providing library preparation and next-generation sequencing services, as well as the Mass Spectrometry Core Facility for providing lipidomics analysis services.

## Data Availability Statement

All data and analyses are available upon request. The RNAseq has been deposited with the Gene Expression Omnibus (Accession: pending), and lipidomics raw files have been deposited with Metabolomics Workbench (Study ID: pending).

