## Supplemental Figures for "Caloric restriction drives age-dependent and integrated remodeling of RNA processing and lipid composition in mouse brain"

**Supplement File Listing**

**Table S1.** Differentially expressed transcripts and corresponding GSEA pathway enrichment results.
**Figure S1.** Rank order plots depicting enriched pathways for 10-month and 20-month CR contrasts.
**Table S2.** Gene expression WGCNA modules and overrepresentation pathway enrichment results.
**Table S3.** Gene expression machinery curated category gene lists.
**Table S4.** Cell-type deconvolution scores.
**Table S5.** Statistically significant alternative splicing events in response to diet.
**Table S6.** Alternative splicing events within the blue and red modules identified via WGCNA on alternative splicing event data.
**Table S7.** Statistically significant lipids in response to diet.
**Table S8.** Individual rCCA sample scores for all comparisons and components.
**Table S9.** Pathways and lipid enrichment analysis results for the transcript-lipid rCCA comparison ranked factors.
**Table S10.** Pathways and alternative splicing event composition for the transcript-splicing rCCA comparison ranked factors.
**Figure S2.** Box plots depicting the individual sample scores for all components and datatypes contributing to the rGCCA analysis.

### Supplemental Figure Legends

**Figure S1. Reactome pathway GSEA enrichments.** GSEA using Reactome pathways of the 10-month and 20-month diet response: normalized enrichment score versus pathway rank, with pathways colored by functional category (transcription and translation; metabolism and mitochondria; RNA processing; neurotransmission and synaptic signaling; signal transduction and cell-fate regulation).

**Figure S2. Per-animal rGCCA sample scores.** Per-animal sample scores from regularized generalized canonical correlation analysis (rGCCA) of the transcript, lipid, and alternative-splicing datasets, for components 1 to 3 (n = 28 animals). Rows are components; columns are datatypes (Transcript, transcript expression; Lipid, lipid abundance; Splicing, alternative-splicing events quantified as percent spliced in). Within each panel, animals are grouped by diet and age (Control or CR at 10, 20, or 30 months; Control n = 5, 4, and 4, CR n = 5, 5, and 5, respectively). Every animal is plotted as a point. Boxes show the median and interquartile range; whiskers extend to the most extreme animal within 1.5 times the interquartile range of the box.
